# Bright Near-infrared Anti-Stokes Fluorescence of ICG under Low Power CW Laser Excitation and its Applications in Bioimaging

**DOI:** 10.1101/2020.06.29.177493

**Authors:** Jing Zhou, Di Wu, Zikang Ye, Dingwei Xue, Mubin He, Jun Qian

**Affiliations:** State Key Laboratory of Modern Optical Instrumentations, Centre for Optical and Electromagnetic Research, College of Optical Science and Engineering, Zhejiang University, Hangzhou, 310058, China; Department of General Surgery, Sir Run-Run Shaw Hospital, School of Medicine, Zhejiang University, Hangzhou, 310016, China; Department of chemistry, Zhejiang University, Hangzhou, 310058, China; Department of Urology, Sir Run-Run Shaw Hospital, School of Medicine, Zhejiang University, Hangzhou, 310016, China

**Keywords:** anti-Stokes fluorescence, FDA-approved molecule, hot-band absorption, near-infrared, high quantum yield

## Abstract

Anti-Stokes fluorescence was observed in ICG, a molecule approved by the FDA for clinical use. The wavelengths of its fluorescence are mainly located in the near-infrared band of 800 nm~900 nm, with a high quantum yield up to 8%. In order to know its generation mechanism, based on multi-photon absorption (MPA) theory, thermally activated delayed fluorescence (TADF) theory and hot band absorption theory, its power dependence, temperature dependence of absorption spectra and fluorescence spectra, and fluorescence lifetime were measured. Its generation mechanism was finally determined to be hot band absorption process. Since ICG showed bright anti-Stokes fluorescence in near-infrared region, which offers substantially longer penetration depth in biological tissues than visible light, excellent photostability and biosafety, we applied it to in vivo imaging and compared it with upconversion nanoparticles (UCNPs). The result is that ICG exhibited much stronger fluorescence than UCNPs, providing more anatomical information of samples. This contributes to a better choice for anti-Stokes fluorescence bioimaging.

## 1 Introduction

Anti-Stokes luminescence is a special optical process, which converts long-wavelength excitation to short-wavelength emission. The generation mechanism of anti-stokes fluorescence can be divided into four classes, multi-photon absorption (MPA) process, upconversion process, thermally activated delayed fluorescence (TADF) process and hot band absorption process^1, 2^.

The MPA process refers to the material simultaneously absorbing multiple long-wavelength excited photons to generate one short-wavelength emitted photon^3^. This process usually occurs in several kind of materials such as dye molecules, AIE nanoparticles, quantum dots, etc^4–6^. The occurrence of MPA emission requires extremely high excitation light energy, so the femtosecond pulse laser is usually used as the excitation light source, which is quite expensive.

During the upconversion process, the upconversion nanoparticle (UCNP) composed of sensitizers and emitters absorbs multiple long-wavelength photons in succession to produce one short-wavelength photon. This process can be achieved by using a continuous wave (CW) laser, which is low cost. Upconversion process is usually divided into lanthanide (Ln)-based upconversion and triplet-triplet annihilation (TTA)-based upconversion^7^. The nanoparticles used for the lanthanide-based upconversion process are mainly composed of lanthanides, the absorption cross section of which is small, resulting in weak upconversion signals^8^. Meanwhile, such inorganic nanoparticles usually exhibit some deficiencies including low solubility and long metabolism time in the organism, which limit their applications in vivo^9, 10^. Another upconversion process, TTA-based upconversion, appears mainly in the nanoparticles containing organometallic complex sensitizers and organic emitters, seldom in the nanoparticles including only organic sensitizers and organic emitters^11^. Such nanoparticles have large absorption coefficients and high quantum efficiency, so their upconversion signals are stronger than those of the lanthanide nanoparticles^12, 13^. However, their photostability is relatively low because of the quenching caused by molecular oxygen^14^.

Thermally activated delayed fluorescence (TADF) is mainly produced by organic molecules. In anti-Stokes luminescence process, the molecule only needs to absorb one long-wavelength photon, and then under thermal activation, it emits one short-wavelength photon. TADF can also be excited by the CW laser.

The generation mechanism of anti-Stokes fluorescence of dye molecules induced by hot band absorption is similar to that of TADF, both of which involve the thermal activation. But in the hot band absorption process, the dye molecule is thermally activated at first, then absorbs one long-wavelength photon, and finally emit one short-wavelength photon. This anti-Stokes fluorescence can be excited by the CW laser. Compared with nanoparticles, the dye molecules can be excreted in a shorter time, imposing less side effects on the living body, which are more suitable for in vivo experiments. To date, most studies have focused on the commercial dyes with high quantum efficiency in the visible range, which are not suitable for in vivo imaging because of severe biological tissue scattering effect. The method of synthesizing molecules that can produce this anti-Stokes fluorescence is not clear, so the research on the hot band absorption materials is still in its infancy.

In our study, an FDA-approved dye molecule^15, 16^, ICG, was found that it can produce bright near-infrared anti-Stokes fluorescence. To explore the potential mechanism about how it produces anti-Stokes fluorescence, a series of confirmatory experiments based on theoretical models were carried out. Moreover, imaging at in vivo level with ICG and comparison with the lanthanide upconversion nanoparticles were further performed to highlight the promising near-infrared anti-Stokes fluorophore. Advantages such as clinical availability, bright near-infrared anti-Stokes fluorescence with reduced scattering effect^17–19^ have suggested ICG’s strong potential for medical application.

## Theoretical models

Unlike the lanthanide nanoparticle or dye nanoparticle, as a dye molecule, ICG’s anti-Stokes fluorescence does not belong to upconversion fluorescence, but belongs to one type of the following three, MPA fluorescence, TADF and hot band absorption fluorescence. The schematic illustration for the three theoretical models has been shown in Fig. 1. There are obvious differences between them, which can be used to determine the generation mechanism of anti-Stokes fluorescence of ICG.

**Fig.1.**
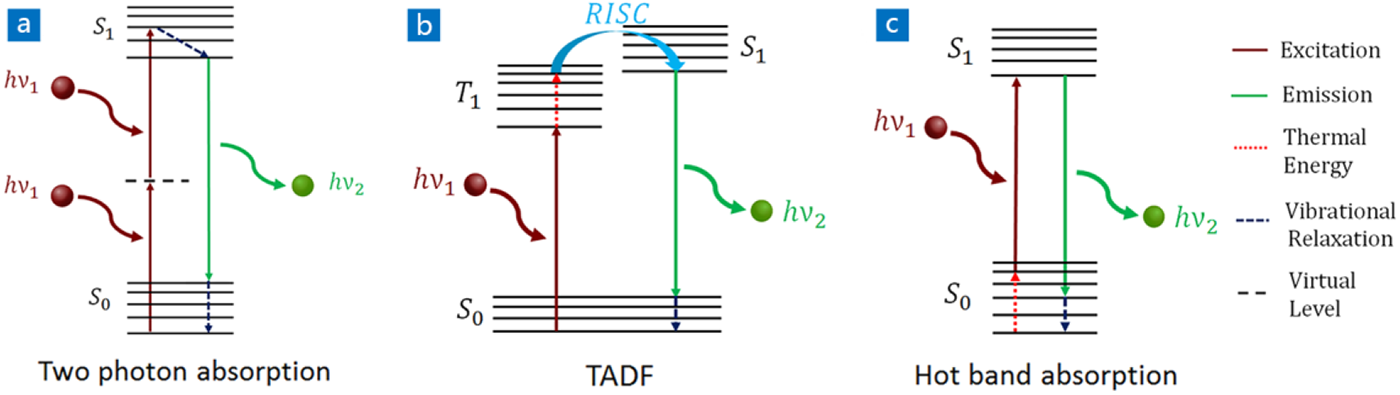
Schematic illustration of different organic molecular anti-Stokes shift luminescence processes.

### MPA process

The schematic of the energy level transition of the molecule in the MPA process has been shown in Fig. 1(a). A molecule in the ground state S_0_ absorbs multiple photons simultaneously to the excited state S_1_ (red solid line arrows). The total energy of the absorbed photons is equal to the molecular transition energy. Then the molecule relaxes to the lowest vibrational energy level of S_1_ (the blue dotted arrow), and eventually transitions back to S_0_, emitting anti-Stokes fluorescence (the green solid arrow).

This is a non-linear optical process. The optical response of a non-linear medium can be determined by the polarization intensity P and the electric field intensity E of the incident light^20^. The formula of the total polarization intensity is shown as follow:

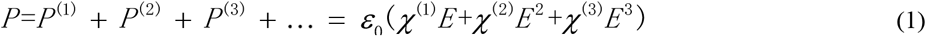

ε_0_ is the vacuum permittivity and χ^(n)^ is the nth-order polarizability of the dielectric material. Here χ^(1)^ is the linear polarization; χ^(2)^ is the second-order polarization; χ^(3)^ is the third-order polarization, χ^(1)^≫χ^(2)^ ≫χ^(3)^. The first term ε_0_χ^(1)^-E corresponds to the absorption, scattering and reflection of light in a linear optical response; the second term ε_0_χ^(2)^ E^2^ reflects second-order nonlinear optical effects such as second harmonic (SHG); the third term ε_0_χ^(3)^ E^3^ describes two-photon and three-photon absorption, third harmonic (THG). When the incident light intensity is weak, the second and subsequent terms can be ignored, and only the first term is retained, which is a linear process. When the incident light intensity is high enough, the higher-order terms can no longer be ignored, so multi-photon absorption process can occur. This is why the MPA process is usually implemented with the femtosecond pulsed laser, rather than the CW laser.

The frequency-upconverted fluorescence intensity Iu is proportional to power n of the excitation intensity Ie, as formula 2 shows,

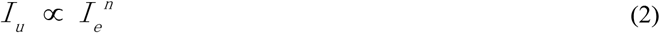

In the case of single-photon absorption process, n=1. If it is a non-linear process, for the two-photon absorption process, n = 2, and for the three-photon absorption process, n = 3.

For the MPA process, the absorption and fluorescence would both be predominantly independent of temperature.

### TADF

In this model, as shown in Fig. 1(b), a molecule in the ground state S_0_ absorbs a long-wavelength photon and directly reaches the triplet state T_1_ (the red solid line arrow), and then undergoes a reverse intersystem crossing (RISC) process from the lowest triplet excited state to the lowest singlet excited state (the blue solid arrow) through thermal activation (the red dotted line arrow), emitting a short-wavelength photon (the green solid arrow). There is also triplet participation in the Stokes luminescence process.

Since the thermal activation process occurs in the triplet state, both anti-Stokes and Stokes fluorescence intensity is positively correlated with temperature. In addition, the molecular absorption coefficients are mainly independent of temperature.

Due to the participation of long-lived triplet states, the lifetime of this type of fluorescence is relatively long (~ μs), which is several orders of magnitude longer than that of singlet fluorescence without triplet participation^21–24^.

### Hot band absorption process

The thermal activation in the hot band absorption process acts on the molecule in the ground state. As shown in Fig. 1(c), a molecule in the ground state with lower energy reaches the ground state with higher energy after thermal activation (the red dotted line arrow), then absorbs a long-wavelength photon to reach the excited state (the red solid line arrow), and finally emits a short-wavelength photon (the green solid arrow)^25–30^.

The occurrence of this type of anti-Stokes fluorescence is closely related to the vibrational energy level of the ground state S_0_. The population of vibrational energy levels in the ground state satisfies the Boltzmann distribution^31^. As formula 3 shows,

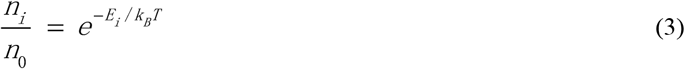

*n*_0_ is the molecular populations of the lowest vibrational energy level in the ground state, while *n_i_* corresponds to the the molecular populations of the higher vibrational energy level *E_i_* in the ground state, kB is the Boltzmann constant, and T is the temperature of the system. The higher the temperature is, the more molecules will be at the higher vibrational energy levels in the ground state, and the fewer molecules will be at the lowest energy level. Molecules in higher ground state vibrational levels will to produce anti-Stokes fluorescence. It can be seen that the absorption coefficients are temperature dependent, therefore, the fluorescence intensity should also be temperature dependent. Besides, the lifetime of anti-Stokes fluorescence is almost the same as that of Stokes fluorescence, both in the order of nanoseconds.

In order to determine which mechanism is responsible for ICG’s anti-Stokes fluorescence, experiments were performed to distinguish these theoretical models.

## Experimental details

### Materials

ICG was purchased from DanDong Pharmaceutical Factory (Liaoning, China). DMSO was purchased from Sinopharm Chemical Reagent Co., Ltd., China. Fresh serum and bile were obtained from rats in our laboratory. The UCNPs (NaYF_4_: Yb^3+^, Tm^3+^) were acquired from Jilin University (Changchun, China). Phosphate buffered saline (PBS) was purchased from Sinopharm Chemical Reagent Co., Ltd., China.

### Animal preparation

Sprague Dawley rats (female, 180g) were used for in vivo experiments. They were provided by the Zhejiang Academy of Medical Sciences and kept at the Experimental Animal Center of Zhejiang University. The room temperature of the rearing environment was maintained at 24 °C with a 12 h light/dark cycle, and the rats were continuously supplied with water and standard laboratory chow. All in vivo experiments were approved by the Institutional Ethical Committee of Animal Experimentation of Zhejiang University and strictly abided by “The National Regulation of China for Care and Use of Laboratory Animals”. The rat was anesthetized at first. Then it was fixed on the imaging platform in the supine position, and shaving and laparotomy were performed to completely expose the imaging area of interest. Finally it was injected with materials through its inferior vena cava.

### Experimental device for measuring absorption spectra, fluorescence spectra and power dependence (915nm CW laser excitation)

The absorption spectra of ICG in DMSO were measured by UV-VIS-NIR spectrophotometer (CARY 5000, Agilent). The fluorescence spectra were acquired from a home-built system based on the PG2000 spectrometer (Ideaoptics Instruments). And the power dependence (915nm CW laser excitation) was derived from the fluorescence spectra excited by different powers.

### Experimental device for measuring lifetime and power dependence (915nm fs pulsed laser excitation)

The femtosecond pulsed laser (PHAROS PH1-10W, LIGHT CONVERSION; repetition rate: 1 MHz; pulse width: 200 fs) beam was introduced into a commercial inverted microscope (IX83, Olympus) as the excitation light. After the laser beam reflected by the dichroic mirror and passing through the objective (PLN20X, Olympus), the sample was excited. By using a turnover mirror, the emitted fluorescence signals passing through the dichroic mirror and filter were either delivered to a spectrometer (Andor, 193i + iXon DU-897U) or detected by the APD (τ-SPAD, PICOQUANT). Fluorescence spectra excited by different powers were measured with the spectrometer and used to analyze power dependence. The computer with integrated time-correlated single photon counting module system (TCSPC, DPC-230 Photon Correlator, Becker & Hickl GmbH) realized the measurement of the fluorescence lifetime of the sample based on the synchronous signals output by the laser and electrical signals from the APD.

### Optical setup for anti-Stokes fluorescence wide-field imaging

The laser beam was coupled to a collimator, and then expanded by a lens with a ground glass sheet to provide a large illumination area. The ground glass sheet was used to eliminate laser speckles. The sample was placed in the position illuminated by the excitation light, and fluorescence signals were captured by a wide spectral response camera (GA1280, 1280 pixels×1024 pixels, TEKWIN SYSTEM, China) after passing through the prime lens (focal length: 35 mm, TEKWIN SYSTEM, China) coated with antireflection (AR) coating at 900-1700 nm and optical filters which were used to filter away the excitation light and extract the fluorescence signals.

## Results and discussion

### Molecular structure, absorption spectra, fluorescence spectra and the anti-Stokes fluorescence image

The molecular structure of the ICG is shown in Fig. 2(a). The black curve in Fig. 2(b) is the absorption spectra (650nm-950nm) of 0.01mg/ml ICG in DMSO. The principle absorption band (λ max = 794 nm) in Fig. 2(b) corresponds to the transition from the lowest vibrational energy level (v_0_) of the ground state (S_0_) to the lowest vibrational energy level (v_0_’) of the excited state (S_1_). The blue edge shoulder at 720 nm is associated with the v_0_ → v_1_’ of the same S_0_ → S_1_ electronic transition, where v_1_’ is a higher vibrational energy level of S_1_^32^. 915nm is at the tail of the absorption spectra. The red curve in Fig. 2(b) is the anti-Stokes fluorescence spectra (< 900nm) of 0.1mg/ml ICG in DMSO under 915nm CW laser excitation. A 900 nm short-pass filter was used. Most of the fluorescence signals are located in the near infrared band of 800nm-900nm. Insert shows the strong anti-Stokes fluorescence from ICG. As previously reported, the full-spectra quantum efficiency of ICG dissolved in DMSO is 13%^33^, so it can be concluded that the quantum efficiency between 800nm-900nm band should reach 8% proportionally.

**Fig.2.**
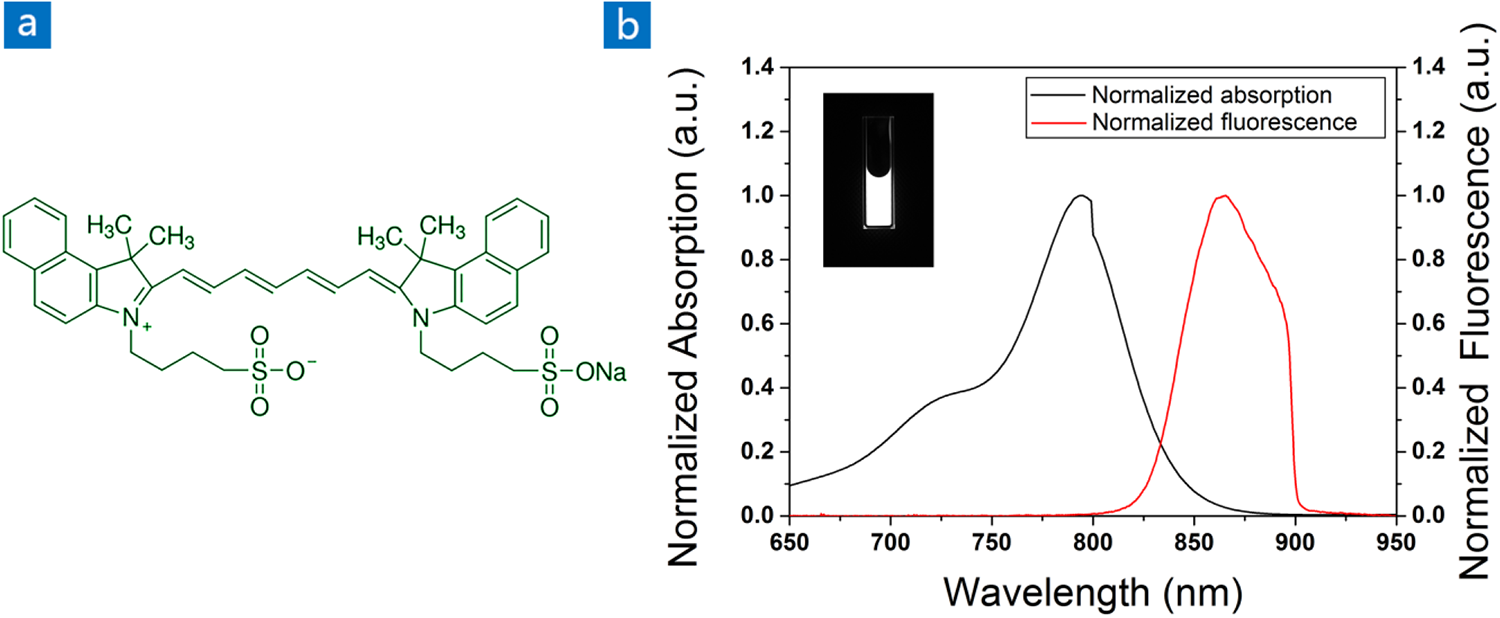
Characterization of ICG molecule. **(a)** Molecular structure of ICG. **(b)** Normalized absorption spectra and normalized fluorescence spectra (900 nm short-pass) of ICG in DMSO. **Insert**: The anti-stokes fluorescence image of ICG (0.1 mg/mL, in DMSO) at the spectral region below 900 nm, under 915 nm CW laser irradiation (26.8 mW/cm^2^), exposure time: 25 ms.

### Power dependence

Since this anti-Stokes fluorescence can be excited by a CW laser, unlike the MPA fluorescence that must be excited by a pulsed laser with high peak power, the first step was to measure the power dependence of ICG in DMSO under 915nm CW laser excitation. The fluorescence spectra of 0.1mg/ml ICG in DMSO excited by different excitation powers were measured with a home-built system. According to the change of the fluorescence intensity at 865nm with the excitation power, a logarithmic plot of power dependence is drawn in Fig. 3. It can be seen that the value of n in formula 2 is 0.99, which is close to 1, indicating a linear anti-Stokes luminescence process, excluding the MPA mechanism.

**Fig.3.**
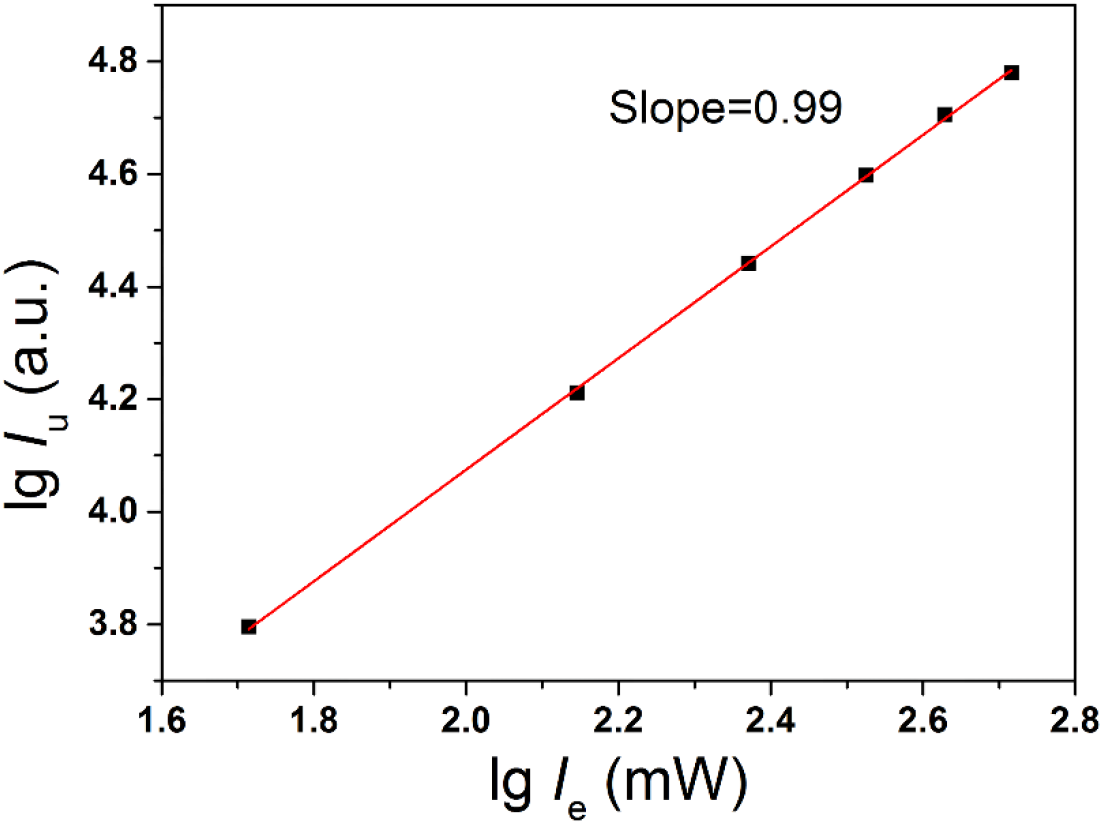
Logarithmic plot of power dependence of fluorescence signal at 865 nm with excitation light (915 nm CW laser), showing a linear dependence of slope 0.99.

### Temperature dependence of the absorption spectra and the fluorescence spectra

The next step was to determine whether the mechanism of anti-Stokes fluorescence is TADF or hot band absorption. The variations of absorption spectra and fluorescence spectra with temperature were measured, as shown in Fig. 4. As the temperature increases, the absorption at long wavelengths increase, while the absorption at short wavelengths decrease. The Boltzmann distribution function shown in formula 3 is a reasonable explanation of this phenomenon. When the temperature rises, the number of molecules at higher vibrational levels in the ground state increases, and the number of molecules at the lowest energy level decreases accordingly. Therefore, with the increasement of the absorption at long wavelengths in Fig 4. (a), both the main band and the higher energy shoulder shown in Fig. 4 (d) decreases at the same time. The anti-Stokes fluorescence spectra excited by 915 nm CW laser in Fig 4. (b) rises correspondingly due to the increased absorption at 915nm as the temperature rises from 303K to 348K, and the Stokes fluorescence spectra excited by 785 nm CW laser in Fig 4. (e) decreases accordingly. Fig 4. (c) and Fig 4. (f) intuitively show that the trends of the absorption at excitation wavelength and the fluorescence intensity excited by it changing with temperature are consistent.

**Fig. 4.**
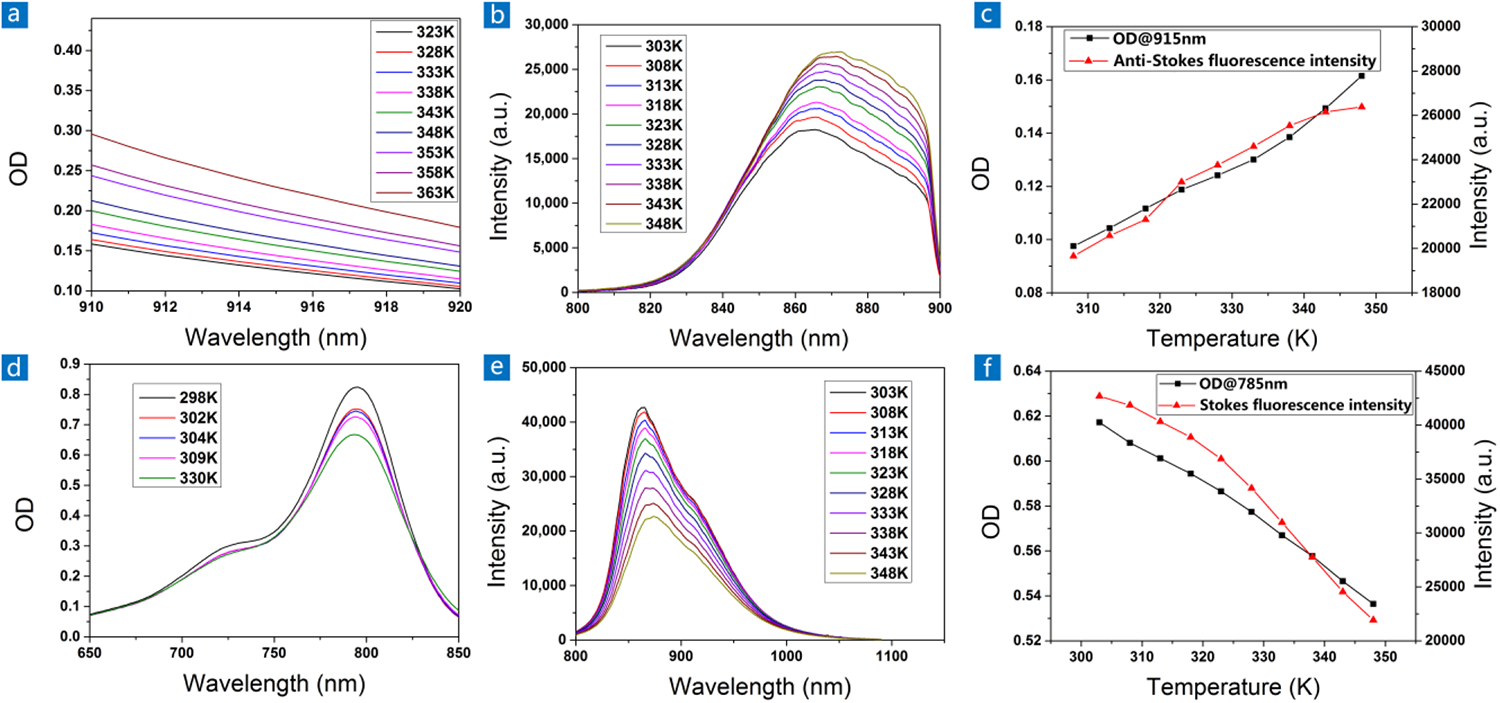
**(a)** Dependence of the absorption spectra (910 nm-920 nm) on temperature. **(b)** The variation of anti-Stokes fluorescence spectra with temperature under the excitation of 915 nm CW laser. **(c)** The black line is the variation of OD at 915 nm with temperature. The red line is the variation of anti-Stokes fluorescence intensity excited by 915 nm CW laser with temperature. **(d)** Dependence of the absorption spectra (650 nm-850 nm) on temperature. **(e)** The variation of Stokes fluorescence spectra with temperature under the excitation of 785 nm CW laser. **(f)** The black line is the variation of OD at 785 nm with temperature. The red line is the variation of Stokes fluorescence intensity excited by 785 nm CW laser with temperature.

Although the temperature dependence of the anti-Stokes fluorescence signals is also consistent with that of TADF, the temperature dependence of the Stokes fluorescence signals and absorption is not. Therefore, the generation mechanism of this anti-Stokes fluorescence is likely to be “hot band absorption”.

### Lifetime of anti-Stokes and Stokes fluorescence

To further determine the mechanism, a time-correlated single-photon counting (TCSPC) equipment was used to measure the lifetime of anti-Stokes and Stokes fluorescence under the femtosecond pulsed laser excitation at 915 nm and 750 nm, respectively. In order to ensure that the measured anti-Stokes fluorescence lifetime did not involve non-linear processes, the appropriate excitation power was explored at first. The power dependence of anti-Stokes fluorescence was measured under the excitation of 915 nm femtosecond pulse laser. As shown in Fig. 5(a), when the power of 915 nm lasers is less than 120 μW after passing through the objective, only linear optical process is involved. So the 915 nm laser power after passing through the objective was set to 4.4 μW, and the 750 nm laser power was set to 49 nW. Then the anti-Stokes and Stokes fluorescence decay curves were measured. The results are shown in Fig. 5(b). The cyan area in Fig. 5(b) is the IRF of this system, whose decay time is 0.25 ns, shorter than that of fluorescence. So this system could be used to measure the fluorescence lifetime of ICG in DMSO. The anti-Stokes and Stokes fluorescence lifetime of 2 mg/ml ICG in DMSO can be obtained as 0.67 ns and 1.21 ns respectively, both on the order of nanoseconds. Therefore, “hot band absorption” should be the mechanism by which this anti-Stokes fluorescence is produced.

**Fig.5.**
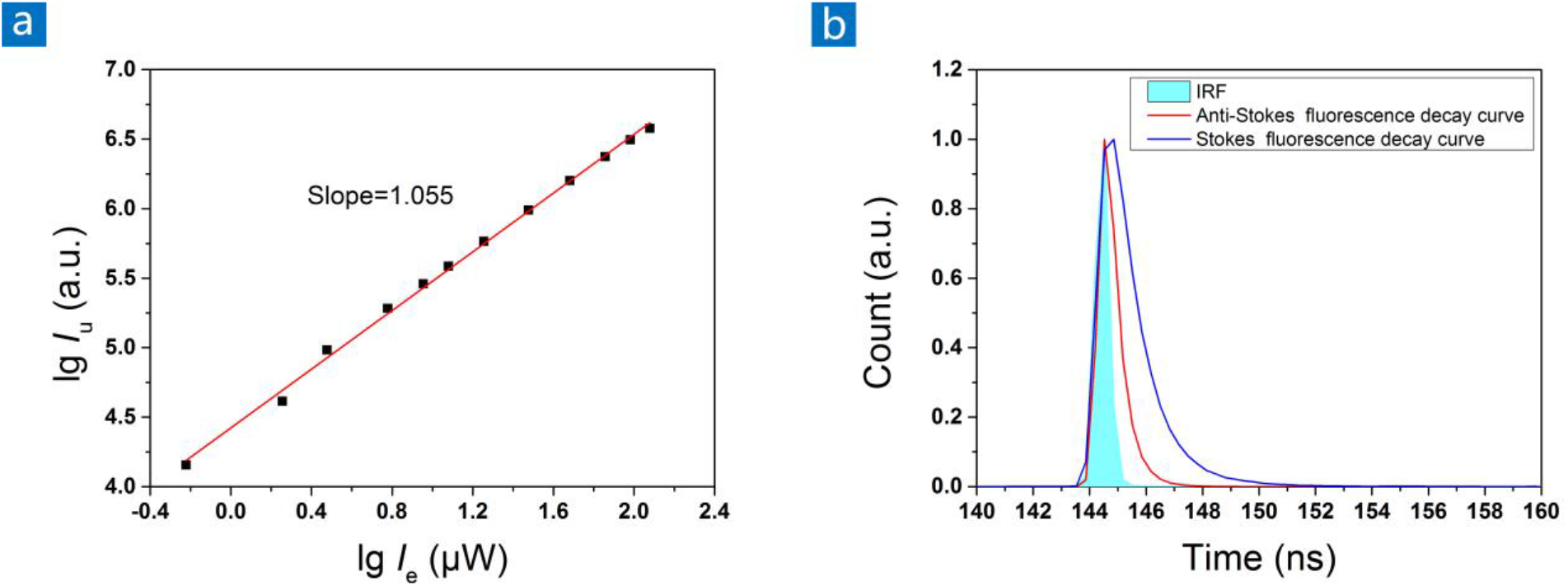
Lifetime of ICG’s anti-Stokes and Stokes fluorescence at a concentration of 2 mg/ml. **(a)** The power dependence of the anti-Stokes fluorescence signal with input power produced by a 915 nm femtosecond pulsed laser. **(b)** The cyan area is the impulse response function (IRF) of the system. The red line is the Anti-Stokes fluorescence decay curve of ICG under 915 nm femtosecond pulsed laser excitation. The blue line is the Stokes fluorescence decay curve of ICG under 750 nm femtosecond pulsed laser excitation.

### Comparison of anti-Stokes fluorescence intensity between ICG and UCNPs in vitro

The generation mechanism of ICG’s anti-Stokes fluorescence has been determined. Considering that it is an FDA-approved clinical fluorescent contrast agent, it was used for bioimaging and compared with UCNPs. UCNPs are a popular class of anti-Stokes luminescent materials that are often used in bioimaging. NaYF_4_: Yb^3+^, Tm^3+^, a type of UCNPs, was selected to do this work. Comparison of ICG and UCNPs on anti-Stoke fluorescence intensity was performed at in vitro level firstly. The anti-Stokes fluorescence spectra of 1 mg/ml NaYF_4_: Yb^3+^, Tm^3+^ in rat serum and bile are shown in Fig. 6(a) and Fig. 6(b). It can be see that the peak fluorescence intensity at 800 nm is only around 2000. However, as shown in Fig. 6(c) and Fig. 6(d), under the conditions of lower excitation power and less integration time, the peak fluorescence intensity at 865 nm of 0.1 mg/ml ICG can reach 20000. Therefore, ICG is much brighter than NaYF_4_: Yb^3+^, Tm^3+^ both in rat serum and bile.

**Fig.6.**
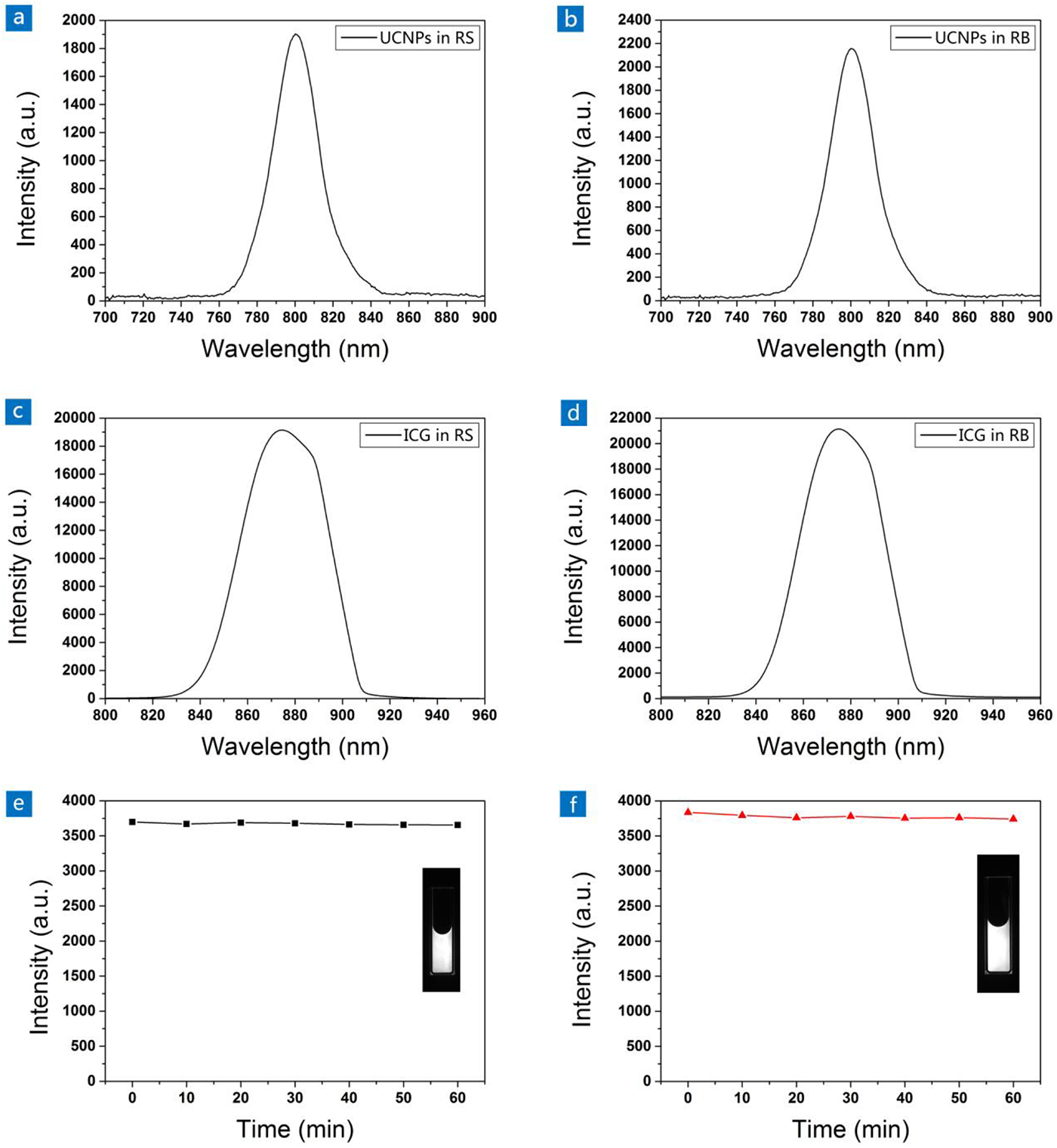
**(a)** Anti-Stokes fluorescence spectra of NaYF_4_: Yb^3+^, Tm^3+^ (1 mg/ml) in rat serum under 980 nm CW laser irradiation (1.8 W/cm^2^). Integration time: 4 s. **(b)** Anti-Stokes fluorescence spectra of NaYF_4_: Yb^3+^, Tm^3+^ (1 mg/ml) in rat bile under 980 nm CW laser irradiation (1.8 W/cm^2^). Integration time: 4 s. **(c)** Anti-Stokes fluorescence spectra of ICG (0.1 mg/ml) in rat serum under 915 nm CW laser irradiation (0.9 W/cm^2^). Integration time: 0.5 s. **(d)** Anti-Stokes fluorescence spectra of ICG (0.1 mg/ml) in rat bile under 915 nm CW laser irradiation (0.9 W/cm^2^). Integration time: 0.5 s. **(e)** Photostability of ICG (0.1mg/ml) in rat serum under 915nm CW laser irradiation (68 mW/cm^2^). **(f)** Photostability of ICG (0.1mg/ml) in rat bile under 915nm CW laser irradiation (68 mW/cm^2^). **Insert**: The anti-stokes fluorescence images of 0.1 mg/ml ICG in rat serum **(e)** and rat bile **(f)** at the spectral region below 900 nm, under 915 nm CW laser irradiation (68 mW/cm^2^), exposure time: 25 ms.

Since dyes often have the disadvantage of poor photostability^34, 35^, photo-bleaching resistance analysis was also performed. As shown in Fig. 6(e) and (f), after the continuous irradiation of 915 nm CW laser whose power (68 mW/cm^2^) was sufficient for in vivo imaging for one hour, ICG showed almost no attenuation of the fluorescence signals, suggesting its excellent photostability under the irradiation of 915 nm CW laser.

### In vivo wide-field imaging

Benefiting from ICG’s bright near-infrared anti-Stokes fluorescence signals and less scattering of near-infrared light in biological tissues which is conducive for biological imaging, ICG was used to visualize blood vessels and biliary tracts in rats. As shown in Fig. 7(c) and Fig. 7(d), the blood vessels in the hind limb of the rat injected with ICG and its biliary tract can be clearly identified, while no fluorescence signal could be detected on the blood vessels and biliary tract of the rat injected with NaYF_4_: Yb^3+^, Tm^3+^ in Fig. 7(a) and Fig. 7(b). Fig. 7(e) and Fig. 7(f) show the FWHM of the hind limb blood vessel and biliary tract of the ICG-injected rat, indicating that the width of the hind limb large blood vessel and biliary tract of the rat are 1 mm and 613 μm respectively. Compared with NaYF_4_: Yb^3+^, Tm^3+^ with the absorption peak at 980nm, another advantage of using ICG’s anti-Stokes fluorescence under excitation at 915 nm for bioimaging is that 980nm is also a water absorption peak, but the absorption of water at 915nm is quite low^36^, which could effectively eliminate the photothermal effect and avoid thermal damage on biological tissues.

**Fig.7.**
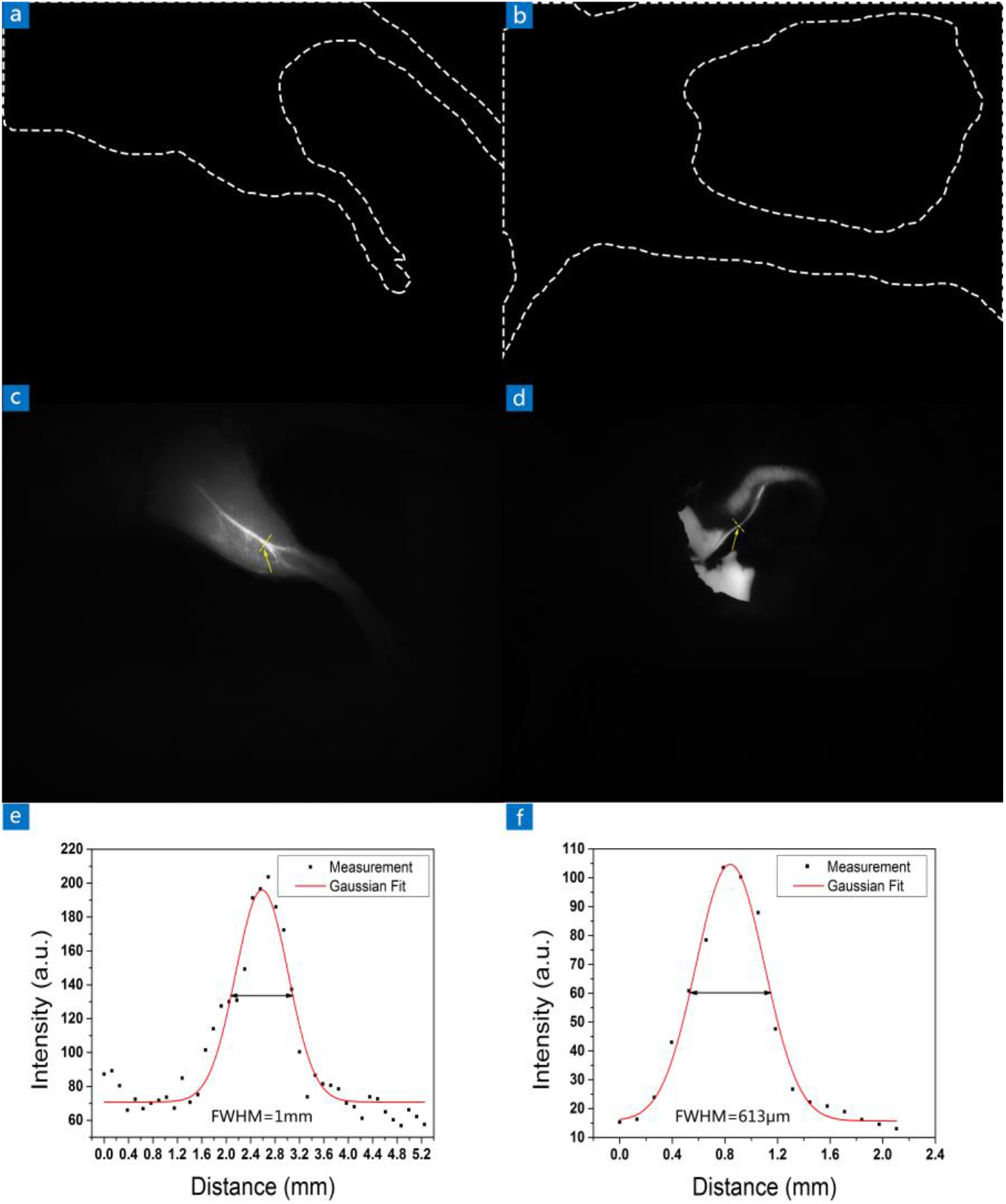
**(a)** In vivo anti-Stokes fluorescence wide-field imaging of blood vessels of a rat after receiving injection of NaYF_4_: Yb^3+^, Tm^3+^ (9.6 mg/ml, 500μl) under 980 nm CW laser irradiation (57 mW/cm^2^). **(b)** In vivo anti-Stokes fluorescence wide-field imaging of the biliary tract of a rat after receiving injection of NaYF_4_: Yb^3+^, Tm^3+^ (9.6 mg/ml, 500μl) under 980 nm CW laser irradiation (57 mW/cm^2^). **(c)** In vivo anti-Stokes fluorescence wide-field imaging of blood vessels of a rat after receiving injection of ICG (1 mg/ml, 200μl) under 915 nm CW laser irradiation (16.5 mW/cm^2^). **(d)** In vivo anti-Stokes fluorescence wide-field imaging of the biliary tract of a rat after receiving injection of ICG (1 mg/ml, 300ul) under 915 nm CW laser irradiation (4.5 mW/cm^2^). Optical filter, 800 nm long-pass and 900 nm short-pass; exposure time, 25 ms. **(e)** The FWHM of the blood vessel pointed by the arrow in **(c)**. **(f)** The FWHM of the biliary tract pointed by the arrow in **(d)**.

## Conclusions

In summary, we found that ICG can produce bright near-infrared anti-Stokes fluorescence, outperforming the conventional UCNPs, and further elucidated the potential luminescence mechanism-“hot band absorption”. Advantages including high quantum efficiency in the near-infrared window, deeper penetrating ability in biological tissues, excellent photostability and FDA-approved suggest ICG’s promising application in bioimaging field. Besides, it also provides a new direction for the design of hot band absorbent materials, which could possibly be excellent candidates for different application areas such as OLED, imaging, sensing and even medical diagnostics and treatment.

## Acknowledgements

This work is supported by the National Natural Science Foundation of China (61975172 and 61735016), and the Zhejiang Provincial Natural Science Foundation of China (LR17F050001).

## Author contributions

All authors commented on the manuscript.

Jun Qian proposed this project and provided guidance and assistance to the project. Jing Zhou is the main executor of the project, exploring the mechanism of ICG’s anti-Stokes fluorescence and using it for bioimaging. Di Wu extracted the serum and bile of rats, and provided assistance for shaving, dissecting and injecting materials to the rats in the in vivo imaging experiments. Zikang Ye helped to measure the fluorescence lifetime of ICG. Dingwei Xue and Mubin He provided some help for the in vivo imaging experiments during the project.

## Authors

Jing Zhou received her Bachelor’s Degree from Department of Optical Engineering of Ocean University of China in 2018. She joined College of Optical Science and Engineering of Zhejiang University for her Ph.D. degree in 2018. She is now engaged in research in the field of biophotonics.

